# HIV-1 evasion of restriction factors: cyclophilin A and cell fusion provide a helping hand

**DOI:** 10.1101/383075

**Authors:** Henry Owen, Alun Vaughan-Jackson, Lea Nussbaum, Jane Vowles, William James, M.D. Moore

## Abstract

Retroviral restriction factors are important regulators of viral infection, targeting vulnerable steps of the virus lifecycle; steps that are also targeted by antiviral drugs. It has become clear that the route of cellular infection can alter the sensitivity of HIV-1 to these agents. Using CRISPR-Cas9 edited pluripotent stem cell-derived macrophages, we have explored the potential of a modified restriction factor (human TRIMCyp) to inhibit HIV-1 replication in both cell free and cell-cell infection models. We show that the expression of TRIMCyp from the endogenous TRIM5α locus potently restricts infection by cell-free HIV-1. Our results also show the importance of the human cyclophilin A-HIV-1 capsid interaction for viral escape from restriction by native human TRIM5α, highlighting the evolutionary interplay between virus and this host restriction factor. However, when co-cultured with infected T cells, stem cell-derived macrophages are primarily infected by fusion between the cells. We have termed infected cells that result from these fusions heterocytia, and show that their formation overcomes multiple restriction factors and the reverse transcriptase inhibitor AZT.

**Importance:** As sentinels of the immune system, macrophages are relatively resistant to infection by pathogens such as HIV-1. However, infected macrophages are found in infected patients and they play key roles in the pathogenesis of the disease as well as being a component of the viral reservoir that must be targeted before treatment can become cure. In this article, we show that some of the mechanisms by which macrophages restrict HIV-1 can be overcome through a recently described cell-cell interaction leading to cell-cell fusion. We also highlight an evolutionary battle between virus and host and show how the virus has co-opted a host protein to protect it from destruction by an antiviral mechanism. These two key findings suggest potential novel treatment strategies that may reduce the viral reservoir and help our natural defences take back control from the virus.

## Introduction

Tissue resident macrophages are important for homeostasis, development and protection from invading pathogens (1). Their role against pathogens includes detection of pathogen-associated molecular patterns (PAMPs) by their numerous pattern recognition receptors (PRRs), inactivation of the pathogen, innate signalling via interferon to bolster the defences of surrounding cells, and antigen presentation to initiate an adaptive immune response. However, being at the first line of defence also makes subversion of macrophages an evolutionarily successful strategy for pathogens. As expressers of both CD4 and CCR5, macrophages stand alongside T cells as a natural host cell for HIV-1, and as a self-renewing population of tissue resident cells they also constitute a long-lived, inaccessible reservoir of virus that hampers viral clearance strategies (2). Consistent with their immune role, macrophages have several antiviral mechanisms to prevent infection by retroviruses such as HIV-1, namely the constitutive expression of retroviral restriction factors, e.g. TRIM5α and SAMHD1 (3).

TRIM5α binds to the viral capsid lattice of an incoming, retroviral core and results in early uncoating, proteasomal degradation of the viral capsid, and ultimately inhibition of reverse transcription; however human TRIM5α is inactive against HIV-1 (4). TRIM5α can be saturated by large doses of virus e.g. during viral dissemination via virological synapses, and can also be circumvented by alterations in the viral capsid gene that prevent TRIM5α binding (5, 6). The compatibility between the viral capsid sequence and the host species TRIM5 locus is a major determinant of successful infection. In fact, converting a single amino acid to the equivalent in rhesus macaque TRIM5α is sufficient to enable human TRIM5α to restrict HIV-1 (7). The same region of capsid that determines the sensitivity of HIV-1 to TRIM5α is also the binding site for the host peptidyl prolyl isomerase enzyme cyclophilin A (CypA) (8). Binding of CypA to an incoming HIV-1 capsid prevents detection of reverse transcription products by macrophages, and the subsequent production of IFN, but conversely binding of CypA aids restriction by TRIM5 of some monkey species (9). This interaction site has been the focus of an evolutionary arms race between the TRIM5 locus and the simian relatives of HIV-1. In several monkey species (e.g. Rhesus macaques and Owl monkeys) multiple retrotransposition events resulted in CypA being inserted into the TRIM5 locus, producing a novel antiviral protein, TRIMCyp (10, 11). Although TRIMCyp does not contain the usual PRYSPRY domain for viral binding, it uses the CypA domain to bind the capsid of sensitive viruses. The species-specific activity of TRIM5 and TRIMCyp suggest that they act as cross-species barriers to retrovirus infection (12, 13), and that in consequence, the TRIM5 locus has great potential to act as a target for gene therapy. To that end, it has been shown that overexpression of macaque TRIM5 variants or TRIMCyp in human cells results in protection from HIV-1 infection (14, 15). However, these approaches fail to take account of the role of endogenous TRIM5α as both a target and signaller of the IFN system (16), nor any potentially adverse consequences of overexpression of this potent antiviral element.

The other major restriction factor expressed by macrophages, SAMHD1, is a deoxynucleotide triphosphohydrolase enzyme (17). The activity of SAMHD1, conversion of dNTPs into deoxynucleosides (dNs), is essential to maintain balanced levels of dNTPs within cells, which is required for genome stability in dividing cells, and is used to reduce the availability of dNTPs to invading pathogens such as HIV and *Leishmania* in non-dividing cells (18-20). In activated T cells, SAMHD1 is both expressed poorly and inactivated by phosphorylation, thus it is unable to inhibit HIV-1 infection (21-23). In contrast, in terminally differentiated macrophages, SAMHD1 is expressed at high levels, is dephosphorylated, and reduces the levels of dNTPs to below the level required by the viral RT enzyme for efficient reverse transcription (19). Although HIV-1 has evolved to be able to replicate, albeit inefficiently, under these conditions, the virus has not evolved a specific counter measure such as VPx of SIV and HIV-2 that targets SAMHD1 for degradation (24), implying that HIV-1 infection is macrophage independent or that the virus has evolved a novel way of bypassing this restriction.

In this report, we use CRISPR/Cas9 to alter the restriction factors TRIM5α and SAMHD1 in order to investigate their impact on the infection of macrophages by HIV-1. By investigating both cell-free and cell-associated infection models we confirm the potency of these factors, but also highlight cell fusion as a mechanism by which they can both be efficiently overcome within tissue macrophages. For this work, we used pluripotent stem cell-derived macrophages (pMac) as a model system for tissue macrophages, as they are genetically tractable at the stem cell stage, karyotypically normal and terminally differentiated, as well as being ontologically, phenotypically and transcriptomically similar to authentic tissue macrophages (25, 26), features not associated with cancerous cell lines or blood monocyte-derived macrophages.

## Results

### Endogenous human TRIMCyp Inhibits HIV-1 Infection

To measure the impact of TRIMCyp in human cells expressed at levels comparable with endogenous TRIM5α, we generated a pluripotent stem cell line containing a knock-in of human CypA into exon 8 of TRIM5α using a double nicking CRISPR-Cas9 approach (Fig 1A) (27). A single clone was identified carrying the CypA insertion in both alleles of TRIM5, as demonstrated by the single peaks of the sequencing trace (Fig 1B) that correspond to the fusion of TRIM5 with CypA at Serine 322 of TRIM5 (28). To observe the effect of human TRIMCyp on HIV-1 infection, the pluripotent stem cells and their differentiated macrophage progeny (pMac) (29) were exposed to increasing doses of a single-cycle GFP-expressing reporter virus, pseudotyped with VSV-G.

**Figure 1.**
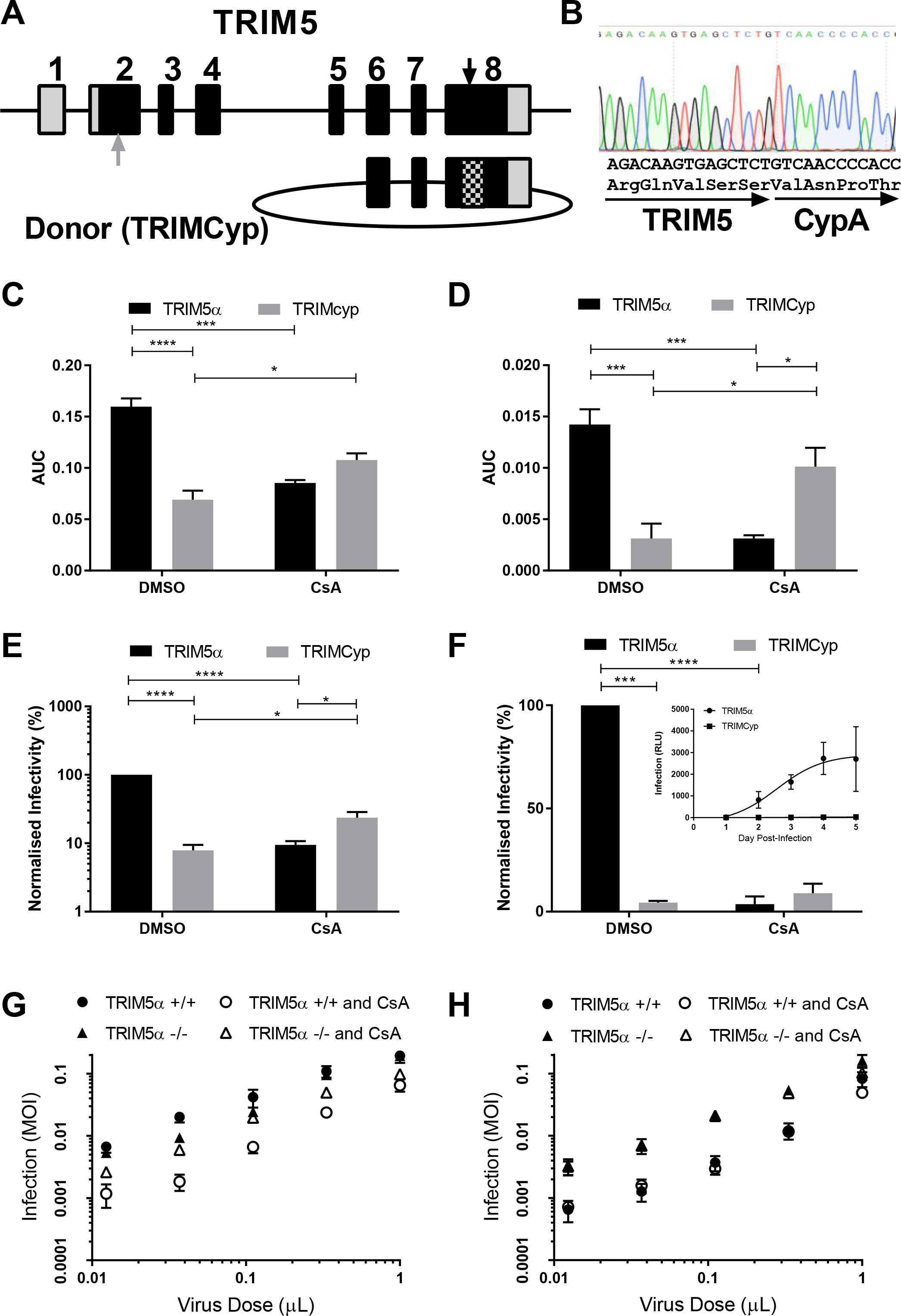
Human TRIMCyp and TRIM5α inhibit HIV-1 of macrophages but are differentially affected by CypA. (A) The TRIM5 locus of a pluripotent stem cell line was targeted by CRISPR-Cas9 nickase at two sites (black arrow) within exon 8 to initiate homologous recombination with a plasmid donor template containing human CypA (hashed box). TRIM5 was also targeted with wildtype CRISPR-Cas9 targeting exon 2 (grey arrow) to produce a knockout phenotype. (B) Sequence trace of the TRIM5 locus at the insertion site of CypA in the identified double knock-in single cell clone. Infection of stem cells (C) and pMacs (D) from an unmodified clone (TRIM5α) and the genetically modified clone (TRIMCyp) with VSV-G-pseudotyped lentiviral vectors, in the presence of 5 μM CsA or DMSO, were quantified by flow cytometry for the reporter eGFP protein. Area under the curve (AUC) analysis of the lower 6 values of one representative serial dilution performed twice (C) or lower 4 values of four (D) independent serial dilutions are shown +/- SEM (*p<0.05, ***<0.001, ****p<0.0001 from a 2-way ANOVA with Turkey’s multiple comparisons test. (E) Infection results (mean +/- SEM) for three different vector constructs all pseudotyped with JRFL envelope, normalized to the infectivity of each vector in DMSO treated wildtype pMacs (TRIM5α) (***p<0.001, ****p<0.0001, for one-sample t-test to hypothetical value of 100, and *p<0.05 for t test). (F) Infectivity of pMacs to replication competent NL-LucR.T2A measured at day 7 post infection in 3–4 independent experiments, plotted as mean +/- standard deviation (***p<0.001, ****p<0.0001 for one-sample t-test to hypothetical value of 100). Inset, a representative growth curve performed in quadruplicate over 5 days showing infection as relative light units (RLU) of luciferase. (G) Serial dilutions of VSV-G pseudotyped eGFP-reporter vector or (H) a P90A capsid mutant vector on pMacs from two wildtype clones (TRIM5α +/+) and two knockout clones (TRIM5α -/-) in the presence of 5 μM CsA or DMSO. The mean +/- SEM of the infectivity, measured as multiplicity of infection (MOI) at each dilution from three independent experiments are shown.

Infectivity in both stem cells and pMacs was reduced by conversion of TRIM5α to TRIMCyp (Fig 1C+D, and FigS1A+S2A). As TRIMCyp is inserted into the TRIM5 locus it can be considered a marker of TRIM5 expression in wild-type cells, thus reduced infectivity of gene-edited stem cells indicates that TRIM5α is normally expressed in pluripotent stem cells, potentially as a mechanism to prevent endogenous and exogenous retroviral infections of the embryo (30). In pMacs, TRIMCyp has a more dramatic impact on infectivity than in stem cells, reducing infection by 10-fold at most viral concentrations (Fig S2A). At higher concentrations of virus (shown in Fig S2A) it is possible to observe the well-established saturation of TRIM-based restriction, such as rhesus TRIM5α and owl monkey TRIMCyp (31). To demonstrate that the reduction in infection, seen in Fig 1C+D and Fig S1A+2A, was due to the modification at the TRIM5α locus and not to any off-target effect, the cells were treated with CsA during infection. CsA prevents CypA from binding the viral capsid and is known to rescue HIV-1 infection in TRIMCyp expressing cells. CsA has also been shown to reduce HIV-1 infectivity in human cells by an incompletely understood mechanism that has been linked to HIV-1 genome detection by cellular DNA detection pathways (9). In agreement with these data, our wild-type cells were more resistant to infection after treatment with CsA (Fig 1C+D, Fig S1B+S2B), whereas treatment of the gene edited stem cells and pMacs with CsA significantly inhibited restriction of HIV-1 by TRIMCyp (Fig 1C+D, Fig S1C+S2C).

### Human cyclophilin A protects HIV-1 from human TRIM5α

Other than overall size of effect, a major difference that is observed between the pluripotent stem cell progenitors and the differentiated pMacs, is a disparity between the unmodified and modified cells when both are treated with CsA (Fig 1C+D). If one compares the results of wild-type macrophages expressing human TRIM5α with those of CsA-treated TRIMCyp-expressing cells (which express CsA-inactivated TRIMCyp, but no human TRIM5α) it would appear that TRIM5α has little or no activity against HIV-1 (Fig S1D+S2D), confirming previously published data (32). However, the comparison of HIV-1-infection in the isogenic TRIM5α and TRIMCyp macrophages both in the presence of CsA, reveals a residual antiviral effect of TRIM5α. If human TRIM5α did not bind HIV-1, and CsA prevented TRIMCyp from binding, then in the presence of CsA both cells should have the same infectivity to HIV-1. However, as can be seen in Fig 1D and Fig S2E, TRIM5α-expressing pMacs treated with CsA are more resistant to HIV-1 than CsA-treated TRIMCyp-expressing pMacs, suggesting an interaction of human TRIM5α with the viral core when CypA binding is prevented. We hypothesise that HIV-1 has evolved to use CypA, to alter its conformation, as an evasion strategy to escape TRIM5α restriction. However, as stem cells did not show this phenotype (Fig 1C and Fig S1E) the impact of TRIM5α on CypA-deficient viruses appears to be cell-type dependent, as has recently been shown for TRIM5α function in Langerhans cells (33), possibly due to reduced effectiveness of downstream effectors in stem cells (34).

As the infection results were obtained using an exogenous reporter (eGFP) virus pseudotyped with VSV-G, an envelope that has been previously been shown to alter susceptibility to TRIM5α restriction (33), we performed the same infection experiments in pMacs using three different vectors that were pseudotyped with JRFL, a cognate macrophage-tropic HIV-1 envelope protein. As the different vectors had different reporter genes (eGFP or luciferase) the results were normalized to the infectivity of wild-type TRIM5α expressing cells. The results confirm those of the VSV-G pseudotyped vectors, with a 10-fold reduction in infectivity in both CsA-treated wild-type cells and in TRIMCyp-expressing cells, but a recovery of infection with CsA treatment of TRIMCyp-expressing cells over that of CsA-treated wild-type cells (Fig 1E).

To observe the impact of TRIMCyp in a more physiological setting, pMacs were exposed to wild-type replication-competent HIV-1 with a luciferase reporter expressed in the Nef open reading frame (NL-LucR.T2A). Tracking the infection over a 5-day period shows a robust growth of HIV-1 in TRIM5α-expressing cells, but not in TRIMCyp-expressing cells (Fig 1F, inset). Although CsA could inhibit replication in TRIM5α-expressing cells, it was not able to significantly alter infectivity of TRIMCyp expressing cells (Fig 1F), possibly due to the multifaceted way CypA is employed by the virus and, therefore, multiple stages of the virus lifecycle that could be affected by CsA (16, 35).

To directly investigate the hypothesis that CypA alters the susceptibility of HIV-1 to human TRIM5α restriction, we generated pMacs knocked out for human TRIM5α using CRISPR/Cas9 (Fig 1A). Exposure of these cells to an HIV-1-based vector clearly confirms previous data showing a lack of robust restriction of HIV-1 by endogenous human TRIM5α (Fig 1G). However, in the presence of CsA the loss of infectivity observed with wild-type cells is abrogated by removal of TRIM5α (Fig 1G), which genetically proves our hypothesis (Fig S2F). To confirm the role of CypA in protection of HIV-1 from TRIM5α, the cells were exposed to the same reporter vector but harbouring the P90A mutation in capsid that prevents CypA binding. In line with the published literature, the P90A mutant is not affected by cell treatment with CsA, but in agreement with our hypothesis, the P90A mutant is more infectious in TRIM5α-knockout pMacs compared with wild-type cells (Fig 1H).

### TRIMCyp is circumvented by cell fusion

As TRIM5α and TRIMCyp restriction can be overcome by saturation with incoming viral cores (31), and it is known that cell-to-cell transmission (e.g. via a virological synapse) can result in large quantities of viral material being transferred to the target cell (36, 37), we set out to establish whether endogenous levels of TRIMCyp were sufficient to protect cells in the more physiological setting of cell-cell contact. To test this, we employed the same experimental set up that was used to demonstrate phagocytosis of infected T cells as a potential route for HIV-1 infection of macrophages (38), with some minor adjustments. As an initial experiment, NL-LucR.T2A infected Jurkat-R5 cells were co-cultured with wild-type or TRIMCyp macrophages for 6 hours and infection of the macrophages was assessed 7 days later (Fig 2A). In contrast to the data obtained by Baxter et al. 2014, we did not see a significant difference between infection levels in the presence or absence of the reverse transcriptase inhibitor AZT, even though in experiments using cell-free virus, AZT reduced infection of the macrophages (Fig 2B). To investigate the reverse transcriptase-independent results further, and to show that the results were not dependent on the genotype of the induced pluripotent stem cell donor, the experiment was repeated using pMacs from three independent donors, using both AZT and the fusion inhibitor, T20 or enfuvirtide (39). Once again, the level of infection was unaffected by AZT, however T20 reduced the level of infection more than 10-fold (Fig 2C), indicating that HIV-1 envelope-driven fusion of the cells was the major cause of the observed infection, a process that appears to be unaffected by TRIMCyp expression.

**Figure 2.**
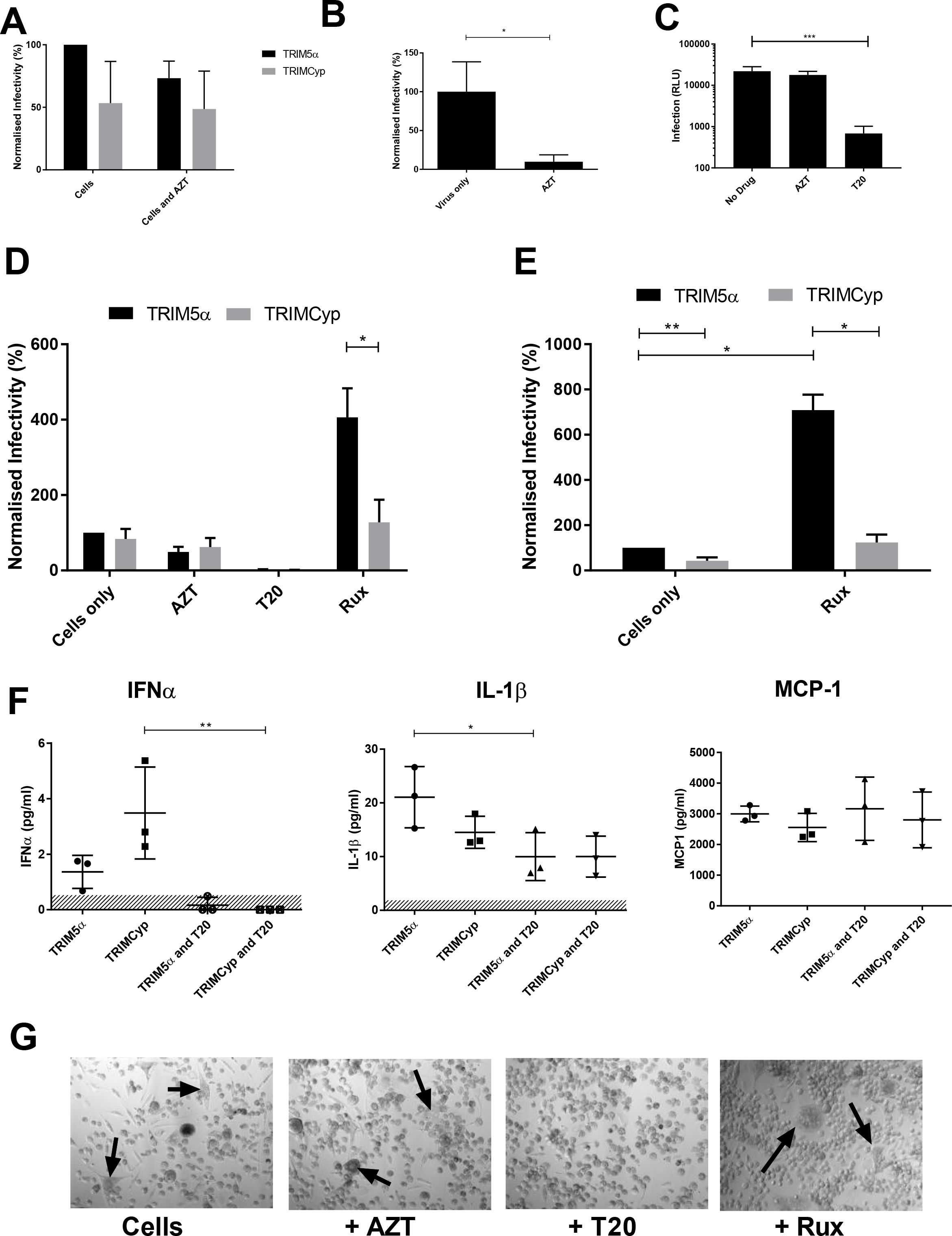
Cell-cell infection of pMacs overcomes TRIMCyp. (A) Macrophages co-cultured with NL-LucR.T2A-infected Jurkat-R5 cells for 6 hours were measured for infection after 7 days, during which etoposide treatment removed residual T cells. Co-cultures were performed in the presence or absence of AZT and are presented as the level of luciferase normalized to the wildtype (TRIM5α) pMacs in the absence of AZT. Results are plotted as mean +/- standard deviation of 3 independent experiments. (B) Effect of AZT on cell free infection of pMacs measured 7 days post infection. Results are plotted as mean +/- standard deviation of 3 independent experiments normalized to the no drug control (*p<0.05, t-test). (C) Effect of anti-HIV drugs AZT and T20 on pMac infection in the cell-cell infection model, co-culturing the cells for 16 hours followed by 4 days etoposide treatment prior to luciferase measurement. Results (raw relative light units or RLU) show the mean +/- standard deviation from three different donor (lines 856–03–04, OX1.19 and HUES2) pMacs (***p<0.001, t-test). (D) Cell-cell infection of pMacs from NL-LucR.T2A infected PBMC-derived CD4+ T cells is enhanced by inhibiting IFN signalling using the JAK1/2 inhibitor Ruxolitinib (Rux). Results are depicted as the mean +/- standard deviation from three different donors normalized to untreated wildtype TRIM5α-expressing macrophages (*p<0.05, t-test with Holm-Sidak method for correction for multiple comparisons). (E) Results from (D) were normalized to the level of infection of each macrophage population in the presence of AZT (*p<0.05, one-sample t-test to hypothetical value of 100, and **p<0.01, t-test). (F) Supernatants from (D) were assayed by Luminex for a variety of cytokines/chemokines, a subset of which are shown. The results are plotted individually with mean +/- standard deviation indicated (**p<0.01, Kruskal-Wallis test, *p<0.05, ANOVA with Sidak’s multiple comparison correction) and the limit of detection is indicated by the hashed area. (G) Representative micrographs of wildtype macrophages on day 7 post coculture with donor derived infected T cells, with black arrows depicting giant syncytial cells.

To test the hypothesis that T cell-macrophage fusion bypasses TRIM-based restriction in a more physiological setting, PBMC-derived CD4+ T cells from three independent donors were infected with NL-LucR.T2A and co-cultured with wild-type or TRIMCyp-expressing pMacs in the presence of AZT, T20 and the IFN-signalling inhibitor Ruxolitinib (Rux). Again, the levels of infection were not significantly altered by the presence of AZT, whereas T20 significantly reduced infection (Fig 2D). Additionally, preventing IFN signalling enhanced infection, particularly in the TRIM5α expressing cells (Fig 2D). By subtracting the background signal of fusion-induced infection (AZT control), it is possible to observe replication of HIV-1 during and/or after the co-culture (Fig 2E), which highlights the significant impact of TRIMCyp expression as well as IFN in this experimental setting. A Luminex^®^ assay on the culture supernatants confirmed the release of IFNα in untreated conditions and an elevation of the proinflammatory cytokine IL-1β, without any impact on the other cytokines/chemokines assayed, e.g. the chemoattractant MCP-1 (Fig 2F).

Upon observing the co-cultures of macrophages and PBMC-derived T cells by light microscopy it became apparent that a major difference between the treatments was the absence of giant syncytial cells in the T20 treated cells (Fig. 2G). It has been known for many years that HIV-1-directed fusion between macrophages can result in giant multinucleated syncytia (40, 41). However, given that the majority of infection in the macrophages was independent of HIV-1 reverse transcription (AZT result in Fig. 2A+D), we hypothesised that the major route of macrophage infection in our experimental setting was T cell-macrophage fusion. To address this, we performed live cell fluorescence microscopy on co-cultures of CellTracker^™^ Orange CMRA-stained pMacs with T cells infected by an eGFP-expressing replication-competent HIV-1. Within 30 minutes, fusion between T cells and macrophages was observed in the co-cultures, with the eGFP-tagged HIV-1 Gag protein being rapidly transferred from the T cell to the macrophage over a period of 5 minutes (Fig 3A and Video S1). This process resulted in macrophages harbouring an infected T cell-derived nucleus in the absence of reverse transcription. The destiny of each virally infected T cell was quantified over a 2-hour period post-co-culture (Fig 3B). The major events included both cell-cell fusion (rapid mixing of the cytoplasmic contents) and cell capture (attachment and colocalisation of a T cell within a macrophage, without cytoplasmic mixing, in a process consistent with phagocytosis of live cells, in Fig S3A, or apoptotic cells in Fig S3B (38). Additional events included rapid loss of GFP signal from the T cell in a process consistent with cell lysis (Fig S3C) and transfer of virus from the T cell to the macrophage in a process consistent with the formation of a virological synapse or plasma membrane transfer between immune cells (Fig S3D) (42, 43). The only event that was modified by antiviral drug treatment was cell fusion by T20, demonstrating that the results for cell-cell infection (Fig 2A+C+D) were the outcome of the formation of a fusion of infected T cells with uninfected macrophages; the product of which we have termed heterocytia, to distinguish them from syncytia, derived from a single cell type, and heterokaryons, derived from genetically different parents.

**Figure 3.**
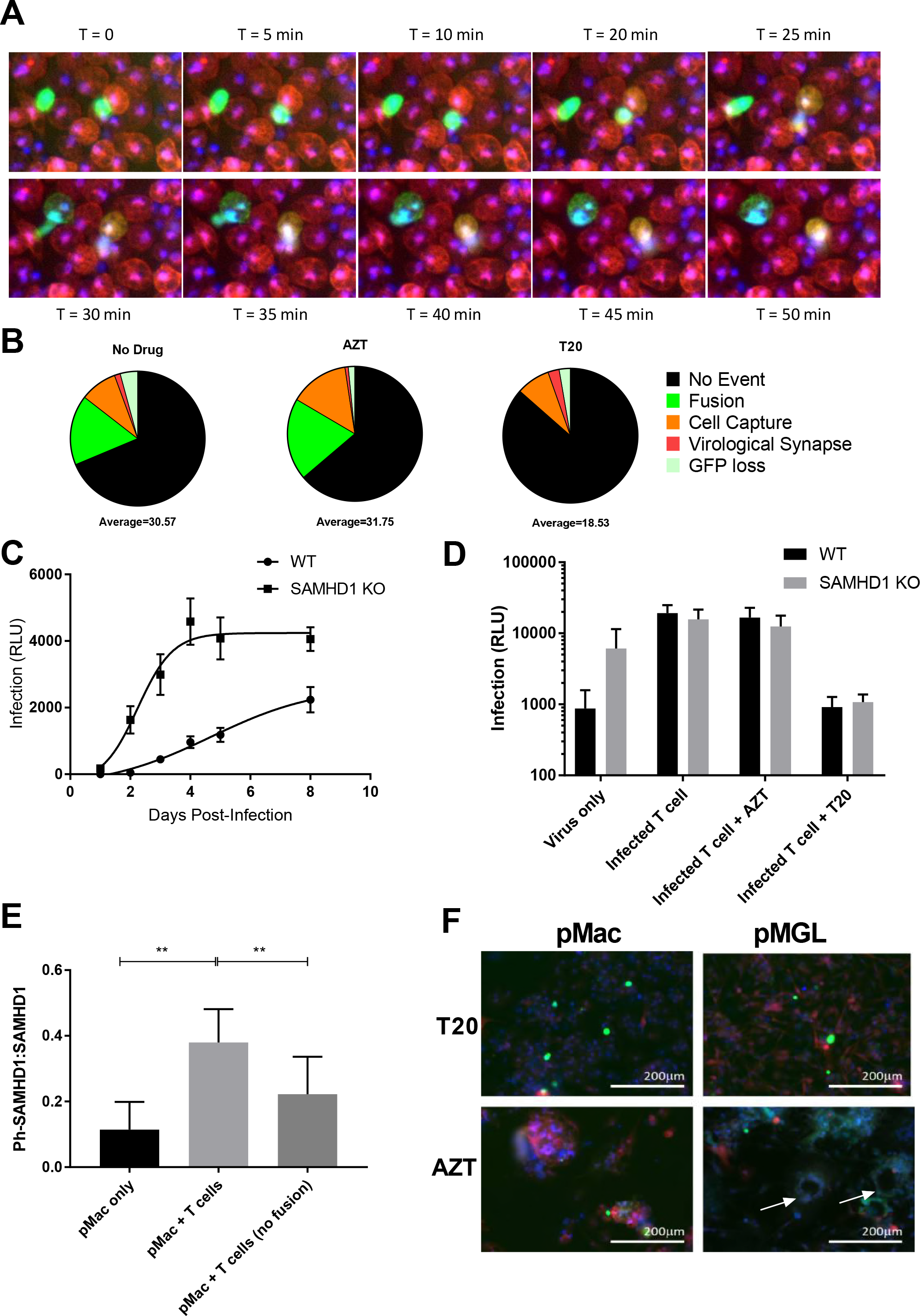
Heterocytium formation results in reverse-transcriptase independent macrophage infection. (A) Representative images of two HIV-1 eGFP-reporter virus infected Jurkat-R5 cells (green) fusing with stem cell-derived macrophages (red). (B) Quantification of cell fates of infected Jurkat-R5 T cells within the co-culture over a 2-hour period. The average number of events from 4 fields of view is also shown. (C) SAMHD1-knockout macrophages show elevated levels and kinetics of infection. The replication of NL-LucR.T2A virus in SAMHD1-knockout macrophages from three independent pMac factories performed in triplicate is shown as mean +/- standard deviation and is compared to that of the parental isogenic wildtype (WT) macrophages. (D) Co-culture of NL-LucR.T2A infected Jurkat-R5 cells with an isogenic pair of macrophages, differing only in the loss of SAMHD1, shows that restriction by SAMHD1 is overcome by cell fusion. Infectivity of the macrophages, in the presence or absence of AZT and T20, was measured 5 days post co-culture with 4 days of etoposide treatment. Results (raw RLU values) are presented as the mean +/- standard deviation of three independent co-culture experiments. (E) Ratio of phosphorylated to total SAMHD1 levels within the co-culture of pMacs and HIV-1 envelope-expressing CEM T cells. The mean +/- standard deviation of 3–5 experiments using two pMac donors are shown, with fusion being prevented by the inhibitors T20 or Tak779, or by use of CEM cells lacking the HIV-1 envelope (**p<0.01, t-test). (F) HIV-1 eGFP-reporter virus infected Jurkat-R5 cells (green), cultured with tRFP-expressing microglial precursors (pMGL, red), both stained with Hoechst 33342 to identify nuclei (blue), form heterocytia with classical giant cell morphology (white arrows) compared with macrophage heterocytia (pMac), in the presence of AZT but not T20.

### Heterocytia formation overcomes SAMHD1 restriction

Given the ability of heterocytia formation to overcome potent TRIMCyp restriction and to result in infected macrophages without reverse transcription, we next investigated if the other major antiretroviral mechanism employed by macrophages, SAMHD1, was also circumvented. We have previously generated macrophages from pluripotent stem cells genetically ablated for the retroviral restriction factor SAMHD1 (44). SAMHD1 knockout macrophages show faster and more efficient infection by HIV-1 when exposed to NL-LucR.T2A virions (Fig 3C). However, when cocultured with NL-LucR.T2A -infected T cells, the isogenic pairs of macrophages, differing only by the presence of SAMHD1, show similar levels of infection (Fig 3D). This infection is unaffected by AZT, but reduced by T20, which demonstrates that cell fusion not only bypasses TRIM5α-mediated restriction but also SAMHD1 restriction, making it independent of the major retroviral restriction factors and, therefore, potentially the most efficient method of macrophage infection.

Given that in dividing cells SAMHD1 is inactivated by cyclin dependent kinase (CDK) phosphorylation at T592 we hypothesized that cell fusion would also alter the regulation of pMac SAMHD1. Within macrophages, SAMHD1 is generally unphosphorylated, and therefore active, whereas in T cell lines such as CEM cells it is not expressed. Therefore, upon fusion between the two cells, any detected increase in phosphorylated SAMHD1 would indicate an interaction between the cells’ proteomes. We co-cultured pMacs from three donors with a CEM cell line constitutively expressing the HIV envelope, for 24 hours before lysing the cells and probing for SAMHD1 phosphorylation by western blot. Quantification of the ratio of phosphorylated SAMHD1 to total SAMHD1 shows that the pMac SAMHD1 is inactivated by fusion with the T cell (Fig 3E and Fig S4E).

### Heterocytia formation occurs in stem cell-derived microglia

To investigate the potential of T cells to fuse with microglia in the brain and thereby generate giant cells, frequently observed in HIV encephalopathy, we employed our recently described and validated pluripotent stem cell model of microglia (26). Differentiating stem cell-derived macrophages in the presence of IL-34 results in microglial progenitor like cells (pMGL) that adopt highly ramified and mobile phenotypes characteristic of microglia in vivo. Addition of T cells infected with an eGFP-expressing replication competent reporter virus to tRFP-expressing pMGLs resulted in not only the heterocytia observed with pMacs, but also giant multinucleated cells with peripherally arranged nuclei that more closely resemble tissue giant cells, e.g. Langhans giant cells, HIV-associated giant cells of the lymphoid Waldeyer’s ring, and HIV encephalitis associated multinucleated cells (Fig 3F) (40, 41).

## Discussion

Macrophages are inherently resistant to HIV-1 infection due to low levels of the viral receptors CD4 and CCR5, high levels of active SAMHD1, and the ability to detect and respond to infections with a robust innate response. Nevertheless, they represent a natural target cell for HIV-1 *in vivo*, contributing to the viral reservoir (45), and are perhaps the most elusive targets for drug treatment, residing in difficult to reach regions such as the CNS, in the form of microglia (46). Most studies into HIV-1 infection of macrophages make use of either monocytic cell lines that have altered cell cycle control and loss of SAMHD1 activity, or blood monocyte-derived macrophages differentiated in vitro under various non-physiological conditions, e.g. high levels of FCS, which is known to alter SAMHD1 activity (47). Given the importance of SAMHD1 in repressing HIV-1 replication, neither of these cellular models is ideal. Moreover, in most tissues monocytes only invade and differentiate into macrophages under inflammatory conditions, so their role in HIV-1 transmission and subsequent pathogenesis is unclear. In contrast, tissue resident macrophages are present within all tissues at steady state, including within the mucosa at the site of infection and, in most tissues, are ontologically distinct from monocytes, being derived from Myb-independent erythro-myeloid progenitors (EMP) of the yolk sac and foetal liver during embryogenesis, not hematopoietic stem cells of the adult bone marrow (2). We have recently shown that pluripotent stem cell-derived macrophages differentiate via an EMP, in a Myb-independent manner (25). Thus, we feel that they are an excellent model of tissue resident macrophages such as microglia, and could help to define the molecular and cellular players during transmission at the mucosa and during neuropathology.

In this work we have used the stem cell-derived model of tissue macrophages to explore the role of TRIM5α on HIV-1 infection. Through genetic manipulation of the human TRIM5α locus to encode the anti-HIV-1 restriction factor TRIMCyp, we have shown that endogenous levels of this protein provide the cells with potent protection from HIV-1 replication. Using CsA to prevent the activity of the CypA component of TRIMCyp, we have also revealed an interesting feature of native human TRIM5α activity. It has long been established that human TRIM5α has little to no activity against HIV-1, however there has been confusion over the impact of CsA on this activity. Initial work suggested that CsA was able to enhance the activity of TRIM5α (48), however the same group and others subsequently presented data that conflicted with this concept (49-51). The current model of CsA action involves destabilization of the viral capsid through loss of CypA binding, resulting in early detection of nascent HIV-1 DNA by cGAS, IFNα production, and suboptimal integration site selection (52). The genetic data presented here shows that TRIM5α does inhibit HIV-1 under conditions in which viral cores lack CypA, either through treatment with CsA or the P90A mutation of Gag. As TRIM5α has been shown to destabilize cores in susceptible virions (53), we hypothesize that without CypA, TRIM5α binds the core, reducing its stability and resulting in detection of the viral DNA. This suggests that HIV-1 evolved to bind CypA as a direct countermeasure to the activity of human TRIM5α, with the added benefit of keeping the virus from detection by the innate immune response. This would suggest that CsA, or one of its non-immunosuppressive variants (54, 55), could be used therapeutically to assist the antiviral effects of endogenous TRIM5α. However, previous attempts using CsA in HIV-1 patients have met with mixed success (56-58). Moreover, as we have identified heterocytium-formation as a potentially significant cell-cell infection pathway, even this approach to activate TRIM5α could be circumvented.

Although infection by cell-free HIV-1 can occur, cell-cell interactions have been shown to enhance transmission between cells. The virological synapse (43), tunnelling nanotubes (59) and phagocytosis of infected T cells (38) have all been observed to enable directed transmission of HIV-1 between cells. However, these all require reverse transcription, a weak link in the viral lifecycle that is targeted by multiple restriction factors (e.g. APOBEC3G/F, SAMHD1, TRIM5) and is a key drug target (e.g. NNRTIs and NRTIs). The data presented here show that cell fusion between T cells and macrophages is an efficient infection process, resulting in an infected heterocytium, that bypasses reverse transcription, thereby avoiding reverse transcription-targeting restriction factors and antiviral drugs.

Syncytium-formation is a fundamental property of mammalian development, being at the heart of trophoblast formation, osteoclast differentiation and muscle fiber generation. However, syncytia are also found in many pathological settings, e.g. Langhans giant cells, giant cells of granulomas and multinucleated giant cells of viral infection. From the perspective of HIV-1 infection, syncytium formation of *in vitro*-infected T cells has historically been linked to CXCR4-tropic viral strains that are also associated with more rapid disease progression (60).

Heterocytia have also been observed within *ex vivo* cultured T cell and dendritic cell co-cultures from tonsil lymph nodes from infected individuals (61). In vivo evidence for the existence of syncytia was for a long time limited to immunohistochemical identification of multinucleated giant cells within the CNS of HIV-associated encephalitis cases (62). More recent studies using humanized mouse models have highlighted a significant amount of small T cell-derived syncytia within the tight confines of lymphoid tissues (63). However, to date these HIV-1-induced syncytia have been reported solely from a phenomenological viewpoint, with little research into their pathophysiological consequences. HIV-1-driven T cell-macrophage fusion has also been previously observed (64, 65), but as low frequency events under certain differentiation conditions that promote the fusogenic properties of macrophages. Our data corroborate this phenomenon and show that it is highly efficient in stem cell-derived macrophages, an authentic cell model of tissue resident macrophages. Such fusion events, *in vivo*, between short-lived infected T cells and long-lived self-renewing tissue macrophages could be of paramount importance in establishing (1) a beachhead at the site of infection, and (2) a viral reservoir in tissues such as the brain as infected T cells transit through the CNS and interact with microglia (66). Additionally, during cross-species transmission events such interspecies heterocytia would allow viral infections to overcome incompatibilities between the virus and innate restriction factors, e.g. TRIM5α. The presence of heterocytia *in vivo* would also provide an alternative explanation for the observation of rearranged TCR DNA within macrophage populations isolated from experimentally infected SIV rhesus macaques (67). Finally, although this study does not address the pathophysiology of these heterocytia during HIV-1 infection, the efficiency of their formation *in vitro* shown in this study, added to *in vivo* observations of syncytia, suggests that they might be a valuable target for treatment. Given that our data demonstrate that the formation of syncytia could be inhibited by the fusion inhibitor T20 and could potentially be limited by IFNα, it is possible that early treatment with these antiviral agents could prevent the neuropathological consequences of infection and reduce the macrophage viral reservoir.

## Materials and Methods

### Cell Culture

Cell culture reagents were sourced from Invitrogen unless otherwise stated. Wild-type human induced pluripotent cell lines SFC840–03–03 (26), SFC856–03–04 (26), OX1-19 (29) and AH016-3 Lenti_IP_RFP (26) and pluripotent stem cell line (HUES2) and its derivatives, have been characterised previously (44). The induced pluripotent stem cell lines were originally derived from healthy donors recruited through the Oxford Parkinson’s Disease Centre having given signed informed consent (Ethics Committee that specifically approved this part of the study, National Health Service, Health Research Authority, NRES Committee South Central, Berkshire, UK, REC 10/H0505/71). All experiments were performed in accordance with UK guidelines and regulations and as set out in the REC. Stem cells were grown in mTeSR^™^1 on Matrigel^®^ (Corning)-coated tissue culture dishes, passaged using TrypLE^™^ and plated with the Rho-kinase inhibitor Y-27632 (10 μM; Abcam). Stem cells were differentiated into pMacs using an established macrophage protocol(29), and where necessary were cultured in microglia induction media as previously described (26). The CCR5-expressing T cell line, Jurkat-R5, was a kind gift from Prof. Q. Sattentau. The HIV-1 envelope-expressing CEM T cell line (CEM-HO-BaL) was engineered using a VSV-G-pseudotyped lentiviral vector co-expressing puromycin and HIV-1 BaL envelope, and a second generation vector (NL4.3R-E-HSA-T2A-Nef) to provide all accessory proteins required for BaL expression. Primary CD4 positive T cells were isolated from donor-derived peripheral blood mononuclear cells (obtained with written informed consent) using the MACS (Miltenyi) CD4+ T cell isolation kit according to the manufactures guidelines and were maintained in RPMI with 10% foetal calf serum and 1% pen/strep and activated using 1 μg/ml PMA and 10 units/ml IL-2 for 24 hrs, 3 days prior to infection with replication competent virus.

### Genome Engineering

TRIMCyp modification: The CRISPR-Cas9 plasmids used in this study were based on the dual guide RNA (gRNA) and Cas9^D10A^-expressing plasmid, pX335-U6-Chimeric_BB-CBh-hSpCas9 (pX335), and its puromycin-resistance gene-expressing derivative, pX462(68) (gifts from Feng Zhang; Addgene plasmids #42335 and #48141). Cloning was performed as previous described(68) using oligonucleotides TRIM1f (CACCGCGAAACCACACGATAATATAT) and TRIM1r (AAACATATATTATCTGTGGTTTCGC) with BbsI digested pX335 to create pX335-Trim51, oligonucleotides TRIM3f (CACCGACAGCACATGAAATGTTGTT) and TRIM3r (AAACAACAACATTTCATGTGCTGTC) with pX462 to create pX462-TRIM3. The donor template was constructed from OX1.19 stem cell cDNA, amplifying three fragments: (1) the 5’ homology arm with primer TRIMcyp1f (TGGTACCGAGCTCGGATCCATGTCAAACACCCAGGAGC) and TRIMcyp1r (TGGGGTTGACAGAGCTCACTTGTCTCTTATCTTC), (2) Cyclophilin A with primer TRIMcyp2f (AGTGAGCTCTGTCAACCCCACCGTGTTC) and TRIMcyp2r (GAGCCCAGGATTATTCGAGTTGTCCACAGTC) and (3) the 3’ homologous arm with primer TRIMcyp3f (ACTCGAAT AAT CCTGGGCTCT CAAAGTATC) and TRIMcyp3r (TGGGCCCTCTAGATGCATGCACACTGCTGGTATATGGAGAG). The three fragments were assembled using the Gibson Assembly^®^ mastermix (NEB), according to the manufacturer’s guidelines, into XhoI and SpeI digested pCR2.1-TOPO (Invitrogen), to generate pTOPO-TRIMCyp.

The pluripotent stem cell line, SFC840–03–03, was Neon^®^ (Invitrogen) transfected with pX335-TRIM1, pX462-TRIM3 and pTOPO-TRIMCyp, as previously (69). Twenty-four hours post-transfection cell were selected for 48 hours in 0.25 μg/ml puromycin. Once recovered the surviving cells were plated on mouse embryonic fibroblasts and single cell cloned, as previously (69). The correct genotype was identified by amplification of the TRIMCyp gene with primers CypAF (CATTGCTGACTGTGGACAACTC) and OutR2 (GCCATTTAAGTATGTTATTCACAG), and was shown to be a homozygous knock-in using primers TRIMsurF (CTGACAGATGTCCGACGCTACT) and TRIMsurR (CGATCAGGACAAATAATCACAGAGA), the product of which was Sanger sequenced using primer TRIMHRMr2 (ACACGTCTACCTCCCAGTAATGTTT)

TRIM5 Knockout: To knockout TRIM5 expression a lentiviral vector (pLentiCRISPRv2.0, gift from Feng Zhang, Addgene plasmid # 52961) was used. Oligos targeting exon 2 of TRIM5α (CACCGTTGATCATTGTGCACGCCA and AAACTGGCGTGCACAATGATCAAC) were annealed, phosphorylated and ligated into BsmBI digested vector as described (70). VSV-G pseudotyped vector was used to transduce SFC840–03–03 cells, which were subsequently selected with 1 μg/ml puromycin for 7 days before single cell cloning on mouse embryonic fibroblasts. Successful knockout clones were identified by Sanger sequencing of the PCR product using primers TRIM5seqF (GTGAAAGCCCTGAGGCATAA) and TRIM5seqR (CCTGCTGAAAGGGGTAATCA) followed by TIDE analysis (71). The two knockout clones used in this work were a homozygous out-of-frame mutant (11 base pair deletion and single base pair insertion) and a heterozygous double out-of-frame mutant (5 base pair deletion in one allele and 4 base pair deletion in the other).

### Lentiviral transduction and Transfection

Lentiviral vectors were generated by PEI-mediated transient transfection of 293T cells using pCMVdeltaR8.2 packaging vector for self-inactivating vectors and either pMD2.G for VSV-G pseudotyped vectors (gifts from Didier Trono; Addgene plasmids #12263 and #12259), or pJRFL for JRFL pseudotyped vectors. The vectors used included pHR’SIN-cPPT-PGK-GIP (a derivative of pHR’SIN-cPPT-EF1-GIP (69), pHIV-HIG (72), pNL4-3.Luc.R-E- (73, 74), pNL4.3R-E- eGFPT2ANef (derived from pNL4-3.Luc.R-E-by replacing the luciferase gene with eGFP). Additionally, pNL4.3R-E-eGFPT2ANef was modified to incorporate the capsid mutant, P90A, using annealed oligos P90Afrd (CAGGGGCTATTGCACCAGGCCAGATGAGAGAACCAAGGGGAAGTGACATAGCAGGAACTA) and P90Arev (CTAGTAGTTCCTGCTATGTCACTTCCCCTTGGTTCTCTCATCTGGCCTGGTGCAATAGCCCCTGCATG).

To replace the fragment released by digestion with SphI and SpeI. Finally, pNL4.3R-E-eGFPT2ANef was modified to include the selectable heat stable antigen (HSA) gene from pHIV-HIG. Where required cells were pretreated with 5 μM CsA or equivalent percentage of DMSO 2 hours prior to infection with viral vectors. Stem cells and pMacs were plated 24 hours or 7 days, respectively, prior to infection in 96 well plates in 100 μl. Infection was carried out by replacing 50 μl of supernatant with 50 μl of three-fold serial dilutions of virus and incubation for 3 days prior to infectivity assay. Fluorescent viruses were quantified by flow cytometry (BD FACSCaliber) and luciferase viruses were quantified by One-Glo^™^ Luciferase assay (Promega) according to manufacturer’s guidelines.

### Replication Competent Assays

The BaL envelope-expressing replication competent viruses either with the reporter renilla luciferase (NL-LucR.T2A (74) or eGFP (NLENG1-BaL-eGFP, a derivative of NLENG1-IRES (75) with the macrophage tropic BaL envelope) in the Nef reading frame, were generated by PEI transfection of 293T cells. Viral infectivity was titred on TZM-bl cells and for infections of Jurkat-R5 cells and primary CD4+ T cells an equivalent MOI of 0.05 was used. Infected Jurkat-R5 cells were used 3 days post-infection whereas the primary CD4+ T cells were used 3–5 days postinfection. For viral growth assays in pMacs the cells were infected at an MOI of 0.02 and the cells were harvested in 24 hour periods and infectivity was quantified using Renilla-Glo^®^ Luciferase assay (Promega) according to the manufacturers protocol. For the cell-cell infectivity assay, NL-LucR.T2A-infected T cells were added to day 7 differentiated pMacs at a ratio of 1:1, with or without pre-treatment of pMacs with AZT (25 μM), T20 (7.5 μg/ml) or ruxolitinib (10 μM). After 6 hours co-culture, the T cells were removed with a single wash in PBS followed by incubation of the cells in macrophage differentiation media supplemented with 25 μM etoposide for 7 days.

### Cytokine release

Supernatant from day 7 co-cultures were assayed by Luminex using an 11-plex custom cytokine panel (Affymetrix) according to the manufactures guidelines, except with the addition of a final fixation with 2% paraformaldehyde for 30 mins at room temperature and two washes before resuspension in reading buffer. Cytokines included IFN-α, IFN-β, IFN-γ, TNF-α, IL-10, IL-1β, IL-6, IL-8, IP-10, MCP-1 and RANTES.

### Imaging of Co-cultures

Day seven differentiated pMacs were stained with 2 μM CellTraker^™^ Orange CMRA before addition of Jurkat-R5 cells, infected 5 days previously with NLENG1-BaL-GFP, at a 2:1 ratio in macrophage differentiation media with 1 μg/ml Hoechst 33342. Cells were imaged on a Zeiss Axiovert 200 microscope with Axiovision MRm camera and Colibri illumination every 5 minutes for 2 hours. Images were processed with Axiovision software (Zeiss) and the fate of all GFP labelled T cells across four fields of view per condition were quantified in a double blind manner.

### Western Blot

CEM-H0-BaL or CEM-H0 T cells were added to 7 day differentiated pMacs at a 3:2 ratio for 16 hours in the presence or absence of 250 nM CCR5 antagonist Tak779 or 7.5 μg/ml T20. Cells lysates (20 μg per lane) were prepared with RIPA buffer (25 mM Tris-HCl pH 7.6, 150 mM NaCl, 1% NP-40, 1% sodium deoxycholate, 0.1% SDS), containing protease inhibitor cocktail (Sigma) and phosphatase inhibitors (sodium othovanadate, sodium fluoride and beta-glycerophosphate), and were separated on NuPAGE Bis-Tris gels (Thermofisher) and transferred onto PVDF membranes. Membranes were probed with mouse anti-SAMHD1 (2D7, Origene, 1:2000), rabbit anti-phT592-SAMHD1 (Cell Signalling Technology, 1:1000) and rabbit anti-GAPDH (Sigma,1:5000) and quantified using the Odyssey far red imager (LiCor).

---

Supplemental Fig 1

Endogenous expression of TRIMCyp is restrictive to HIV-1 in pluripotent stem cells. Serial dilutions of a VSV-G pseudotyped lentiviral vector were used to infect pluripotent stem cells of the wildtype clone (TRIM5α) and the genetically modified line (TRIMCyp). The multiplicity of infection (MOI) at different dilutions of virus, as measured by flow cytometry for the reporter gene eGFP, are shown as the mean +/- standard deviation of a representative experiment performed in triplicate. (A-E) depict different combinations of cell types and treatments (CsA at 5μM) for clarity, all of which is summarized in Fig. 1C. (F) Schematic representation of the interplay between HIV-1 core (image adapted from (76), the PRYSPRY domain of TRIM5α (red), the Cyp domain of TRIMCyp (blue) and the host cyclophilin A (blue ovals) in the presence and absence of cyclosporin A (CsA).

Supplemental Fig 2

Endogenous expression of TRIMCyp is restrictive to HIV-1 in stem cell-derived macrophages. Serial dilutions of a VSV-G-pseudotyped eGFP-expressing lentiviral vector were used to infect pMacs derived from the wildtype stem cell clone (TRIM5α) and the genetically modified line (TRIMCyp). The multiplicity of infection (MOI) at different dilutions of virus, as measured by flow cytometry for the reporter gene eGFP, are shown as the mean +/-standard deviation of four independent experiments. (A-E) depict different combinations of cell types and treatments (CsA at 5μM) for clarity, all of which is summerised in Fig. 1D.

Supplemental Fig 3

Cell-cell interactions in co-cultures of pMacs and HIV-1 infected T cells. Jurkat-R5 cells, infected with a replication-competent eGFP-reporter virus, were co-cultured with CellTracker Red labeled pMacs and imaged over time. Representative images are shown depicting (A) cell capture (phagocytosis), (B) apoptosis followed by phagocytosis, (C) GFP loss (cell lysis), (D) viral transfer (highlighted with white arrow). (E) Effect of fusion on SAMHD1 phosphorylation. A representative western blot shows the level of total (SAMHD1) and T592-phosphorylated (P-SAMHD1) SAMHD1 as well as the loading control (GAPDH). Macrophages alone (pMac only), CEM T cells alone (CEM only), and co-cultured cells incapable of fusing either through lack of HIV-1 envelope (pMac+CEM) or addition of a fusion inhibitor (pMac+CEMH0-BaL+Tak779) were compared to co-cultures allowed to fuse (pMac+CEMH0-BaL).

Supplemental Fig 4

Quality control of the TRIMCyp stem cells and pMacs. (A) SNP analysis (OmniExpress24 chip) of the parental line, wildtype clone (TRIM5α) and modified line (TRIMCyp) are shown, analysed using KaryoStudio (Illumina) to detect copy number variations across the genomes. Duplications are shown in green, deletions in red and loss of heterozygosity in grey. (B) Flow cytometry analysis of antibody staining for key pluripotent stem cell markers, TRA-1–60 and NANOG (dark grey), on the stem cell clones of wildtype and genetically modified cell lines versus their respective isotype controls (light grey). Stem cells were fixed in 2% paraformaldehyde, followed by re-suspension in ice-cold 100% methanol before washing in FACS buffer (PBS with 10 μg/ml IgG from human serum, and 2.5% FBS). Antibodies used included anti-Tra-1–60-Alexa^488^ (Biolegend), anti-NANOG-Alexa^647^ (Cell Signaling) and isotope control antibodies IgM-Alexa^488^ (Biolegend) and IgG-Alexa^647^ (Cell Signaling). (C) Flow cytometry analysis of antibody staining (dark grey) for the myeloid specific lineage marker CD14 and CD68, versus their isotype controls (light grey) on pMacs differentiated from the wildtype and modified cell lines. Antibodies used included anti-CD14-FITC, anti CD68-FITC and their respective isotype controls (all from Immunotools).

Supplemental Video 1

Macrophage infection though cell-cell fusion. HIV-1 eGFP-reporter virus infected Jurkat-R5 cells (green) were cultured with CellTracker Red labelled stem cell-derived macrophages (red), in the presence of Hoechst 33342 to identify nuclei and imaged every 5 minutes. The video shown is a representative video of those used to generated data for Fig 3A+B+S3. Multiple fusion events can be observed as eGFP expressed from the T cells is rapidly transferred to the macrophages.

## Acknowledgments

This publication arises from research funded by the John Fell Oxford University Press (OUP) Research Fund. The pluripotent stem cell lines used in this study was originally generated from donor samples supplied by the Oxford Parkinson’s Disease Centre (OPDC) study (funded by the Monument Trust Discovery Award from Parkinson’s UK, a charity registered in England and Wales (2581970) and in Scotland (SC037554), with the support of the National Institute for Health Research (NIHR) Oxford Biomedical Research Centre based at Oxford University Hospitals NHS Trust and University of Oxford, and the NIHR Comprehensive Local Research Network), and was reprogrammed within StemBANCC, (supported by the Innovative Medicines Initiative Joint Undertaking under grant agreement number 115439, resources of which are composed of financial contribution from the European Union’s Seventh Framework Program (FP7/2007e2013) and EFPIA companies’ in kind contribution). AV-J and LN are supported by the Infection, Immunology and Translation Medicine Wellcome Trust-funded DTC (grant numbers 203805/Z/16/Z and 108869/Z/15/Z, respectively). The funders had no role in study design, data collection and interpretation, or the decision to submit the work for publication.

## Author Contributions

MDM designed, carried out, experiments, analysed data and prepared the manuscript. HO, AV-J, LN, JV carried out experiments. WJ and AVJ analysed data and prepared the manuscript.

